# Vehiculation and functional delivery of lipophilic therapeutics and antibiotics via pulmonary surfactant in a lung-on-chip model

**DOI:** 10.64898/2025.12.18.695116

**Authors:** Cristina Garcia-Mouton, Richa Mishra, Kunal Sharma, Theo Nass, Michelle Sommer, Jesus Perez-Gil, Vivek V. Thacker

**Author notes:** Senior authors, these authors co-supervised the work (Jesus Perez-Gil, Vivek V. Thacker).

## Abstract

Pulmonary surfactant forms a dynamic proteolipid thin-film at the alveolar air-liquid interface. It is both a barrier to but also a “last-mile” carrier for inhaled particulate matter to the distal lung. The role of vehiculation in pulmonary delivery of inhaled therapeutics remains poorly understood, as it cannot be easily studied in animal models, and is not recapitulated in liquid cell culture. Here we adapted a thin-film bridge (TFB) delivery method from vehiculation experiments in acellular models to a lung-on-chip (LoC) platform under breathing-like stretch. Using confocal live-imaging, we demonstrate vehiculation and cellular uptake of fluorescently-labelled cargoes such as Tacrolimus or Beclomethasone. We systematically compared TFB vehiculation to existing in vitro delivery approaches. TFB vehiculation promoted sustained retention of functionally effective Tacrolimus, strong co-localization with surfactant lipids, and accumulation in macrophages in human LoCs reconstituted with in vitro–differentiated alveolar macrophage-like cells. This contrasted with the unphysiological dominant uptake of formulations in liquid by alveolar epithelial cells. Finally, in proof-of-concept experiments, TFB-vehiculated bedaquiline effectively supressed subsequent growth of *Mycobacterium tuberculosis* in a prophylactic manner. Overall, our work demonstrates an approach for studying drug vehiculation in models of the alveolar interface in vitro, and the feasibility of surfactant-containing therapeutic formulations for direct and effective pulmonary delivery.

## Introduction

In the vast surface area of the distal lung, the alveolar-blood interface consists of alveolar epithelial cells and lung microvascular endothelial cells closely apposed to each other, providing the thin barrier needed for gas exchange ^1^. The air-liquid interface is a defining physiological feature of this environment. In healthy individuals, the distal airways are lined by a continuous film of pulmonary surfactant that coats the very thin aqueous layer that provides hydration and protection to the respiratory epithelium ^2^. This film consists primarily of dipalmitoylphosphatidylcholine (DPPC), other phosphatidylcholines (PC), smaller amounts of other phospholipids including unsaturated phospholipids, cholesterol and four proteins – the hydrophobic SP-B and SP-C proteins that are indispensable for surfactant function, and hydrophilic SP-A and SP-D ^3,4^. Together this proteolipid complex: (1) reduces to a minimum the surface tension at the interface, enabling breathing with little energy costs ^5,6^, (2) constitutes the mucosal immune component of the lung, creating a cross-talk between epithelial cells and alveolar macrophages through a cycle of secretion and recycling ^4,7–9^, (3) establishes a barrier against exposure of lung epithelial cells and macrophages to inhaled substances and yet paradoxically (4) is also the last-mile delivery mechanism for inhaled substances/particles to the distal lung ^10,11^. This complex set of functions results from the close interplay between surfactant composition, structure and function, and the airflow dynamics in the lung, which dictates that only < 5 µm diameter aerosols can be directly delivered to alveoli. All particles of sizes above this cutoff impinge upon the upper airway and respiratory bronchioles and thus can only be transported to the alveoli by vehiculation.

Overall, the *in situ* function of pulmonary surfactant, particularly in the context of immune function or as a protective barrier remain poorly understood, as perturbations to this system in animal models are often lethal, its biophysical form and the air-liquid interface are tightly connected and the air-liquid interface in the distal lung can only be accessed via specialized intravital microscopy. These factors have stymied the overall development of the pulmonary delivery route for therapeutics, even though the large alveolar-blood interface provides a promising route for drug delivery with increased efficacy and reduced toxicity – either targeted directly to the airways or for systemic vascular dissemination.

In recent years, more complex in vitro microphysiological models have provided the capabilities to address some of these limitations. Models including Transwells and organ-chips have provided in vitro platforms to study the biology of respiratory cells and in particular epithelial cells at the air-liquid interface, both in the context of homeostasis, but also for host-pathogen interactions in viral ^12,13^, bacterial ^14^ and fungal infections ^15^, sterile inflammation or regeneration ^16^. Lung-on-chip models typically include co-culture of at least epithelial and endothelial cells, often on either side of a membrane that can be stretched in one or more dimensions ^17–19^. In certain systems, the vascular cells are also exposed to flow. Yet despite this increased complexity, the biophysical recreation of pulmonary surfactant function in the airway channel of lung-on-chip models is underdeveloped and understudied. In these systems, delivery of therapeutics, foreign, or infectious agents to the airway channel continues to be via liquid inoculation without the involvement of any pulmonary surfactant component, no different from that in standard cell culture. Several crucial aspects of how the lung barrier interacts with the environment, and with particulate matter in the airways are thus not captured.

Here we adopt techniques used in acellular experiments to develop and demonstrate methodologies for the vehiculation of exogenously provided pulmonary surfactant formulations directly onto the alveolar epithelial cells in a commercially available lung-on-chip system. Live-cell imaging of fluorescently tagged lipids shows that vehiculation at the air-liquid interface results in a more physiologically relevant uptake of surfactant lipids. Using a fluorescently labelled version of the lipophilic immunosuppressive small molecule Tacrolimus as a cargo, we demonstrate feasibility for delivery via vehiculation and functional effectiveness in reducing inflammatory responses to sterile inflammation. Finally, in proof-of-concept experiments, we also demonstrate that the delivery of a lipophilic antibiotic like bedaquiline via vehiculation has a prophylactic and long-lasting antimicrobial effect upon subsequent bacterial infection. Our methodology could be extended to several available organ-chip models.

## Main

We established primary murine lung-on-chip models as previously described ^20^, using alveolar epithelial cells and lung microvascular endothelial cells from C57BL/6 mice at low passage number, and bone-marrow derived macrophages from C57BL/6-Tg(CAG-EGFP)131Osb/LeySopJ mice that constitutively express GFP (Fig. 1a). This allowed us to identify macrophages in co-culture with epithelial cells at the air-liquid interface using confocal imaging. Tacrolimus is an anti-inflammatory immunosuppressive drug that is often prescribed to patients receiving organ transplant ^21^. Its lipophilic nature makes delivery to pulmonary cells challenging, making it an attractive molecule for airway delivery ^22^. We first evaluated the uptake of Tacrolimus conjugated to NileBlue (TacNB) in the murine LoC, imaging the epithelial face 6 hours post-incubation with a 2% weight-by-volume (w/v) solution of TacNB (Fig. 1b). A very low intensity was observed, with TacNB only in a small minority of epithelial cells (Fig. 1c, left). Next, we evaluated if co-association with pulmonary surfactant would alter this outcome. We dried organic extracts of porcine pulmonary surfactant (OE) generated in the Perez-Gil lab with a 1% mol/mol amount of Rhodamine-conjugate 1,2--dioleoyl-phosphoethanolamine (RhodDOPE) together with 2% w/w of TacNB ^10^. This lipid film was then gently hydrated under periodic shaking in PBS at 45°C, above the transition temperature of pulmonary surfactant phospholipids, to ensure uniform incorporation and dispersion of TacNB and RhodDOPE into the unilamellar and multi-lamellar surfactant assemblies that form in liquid solution (Fig. 1b). Under similar experimental conditions, this led to a significant increase in the intracellular levels of TacNB (Fig. 1c, right, 1d). Enhanced uptake by alveolar epithelial cells was also observed when beclomethasone dipropionate also conjugated with Nile Blue (BDP-NB) was used as the cargo, showing that these observations were not specific to TacNB (Fig. S1a). To resolve whether this increase in uptake required the presence of hydrophobic surfactant proteins, we generated multi-lamellar suspensions of DPPC:POPG at a ratio of 7:3 in which TacNB was incorporated as described above. Increased uptake and alveolar epithelial cell localisation of TacNB were also observed the multi-lamellar suspension was inoculated into the epithelial channel of the LoC (Fig. S1b). Thus, we conclude that incorporation into pulmonary surfactant organic extracts substantially increases delivery to epithelial cells, likely by enabling uptake and retention through endocytosis, and avoiding non-specific uptake and efflux.

**Fig. 1.**
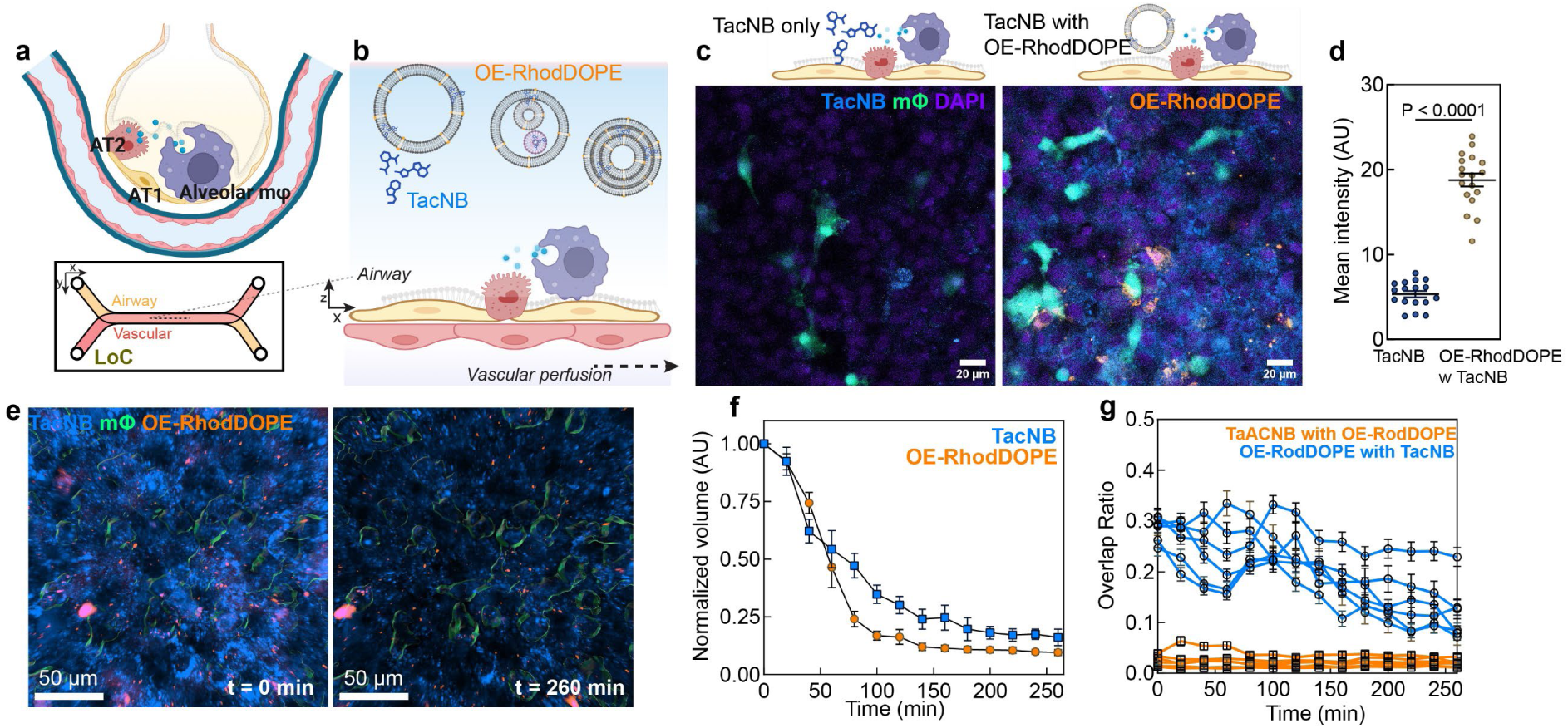
Delivery of Tacrolimus in liquid formulations with pulmonary surfactant enhances uptake by alveolar epithelial cells. (**a**) Schematic of alveolus and lung-on-chip model. (**b**) Schematic of delivery of Tacrolimus-NileBlue (TacNB) alone or TacNB with Organic Extract containing 2% RhodDOPE (OE-RhodDOPE) by addition of liquid to the epithelial channel. (**c**) Representative confocal image of the epithelial face of a murine LoC after inoculation with TacNB alone or TacNB with OE-RhodDOPE and mean intensity (**d**). TacNB – blue, macrophages – green, OE-RhodDOPE – orange. (**e**) Representative 3D views from live-cell confocal imaging of the epithelial face of a murine LoC, macrophages are shown as surfaces. (**f**) Volume of TacNB and OE-RhodDOPE on the epithelial face normalized to the volume at t=0. Data from *n=6* fields of view. (**g**) Overlap ratio between TacNB and OE-RhodDOPE (orange) and the converse ratio between OE-RhodDOPE with TacNB (blue), data from *n=6* fields of view. Data shown is mean ± s.e.m., P-values determined by a two-tailed Mann-Whitney test in (**d**).

To further characterize the dynamics of this process, we performed time-lapse imaging for TacNB delivered in pulmonary surfactant formulations, imaging the epithelial face of the LoC at regular intervals after the removal of the formulation. The use of 3D z-stacks allowed for identification of intracellular components (Fig. S2a). Next, we performed confocal time-lapse imaging, commencing immediately after withdrawal of the formulations from the airway channel. Even at the early timepoints, we observed that both the OE-RhodDOPE and TacNB were taken up predominantly in the alveolar epithelial cells (Fig. 1e, left), and the total volume of both components declined rapidly within 90-120 minutes of culture at the air-liquid interface (Fig.1e, right, 1f). Using Imaris, we segmented areas of TacNB, OE-RhodDOPE and macrophages as surfaces, and quantified the overlap ratio between surfaces (Fig.1g, S2b,c). There was a poor co-localisation of OE-RhodDOPE and TacNB, reflected in a very low overlap ratio between TacNB and OE-RhodDOPE, consistent with the large component of free TacNB in the epithelial layer (Fig. 1g). The proportion of OE-RhodDOPE coinciding with TacNB was also low and decreased over time (Fig. 1g) and both components showed low uptake in macrophages (Fig. S2b,c). Thus, the pattern of uptake and delivery was unphysiological in several respects, most notably because aerosolized particles are primarily observed in alveolar macrophages *in vivo*.

We therefore devised a new strategy for airway delivery that would mimic the vehiculation-based delivery expected *in vivo*, by adapting techniques developed in acellular systems. In previous studies, the Perez-Gil lab has developed a customized double balance setup where a thin paper bridge joins a donor trough (representing the upper airway) with a recipient trough (representing the distal airway) ^10,23^(Fig. 2a). This was replicated with the LoC, where the solution of OE-TacNB vesicles was carefully pipetted on the surface of the donor receptacle, which was then connected to the input of the LoC device via a thin wedge of Whatman paper ^10^ wetted with PBS (Fig. 2b). Prior to the start of transfer, the airway surface of the LoC was washed with PBS, and the LoC was maintained under breathing conditions throughout, which was necessary for effective transfer. At the end of the 30-minute period, the bridge was removed, and the LoC maintained at air-liquid interface with vascular perfusion (Fig. 2c). We term this method as thin-film bridge (TFB) vehiculation.

**Fig. 2.**
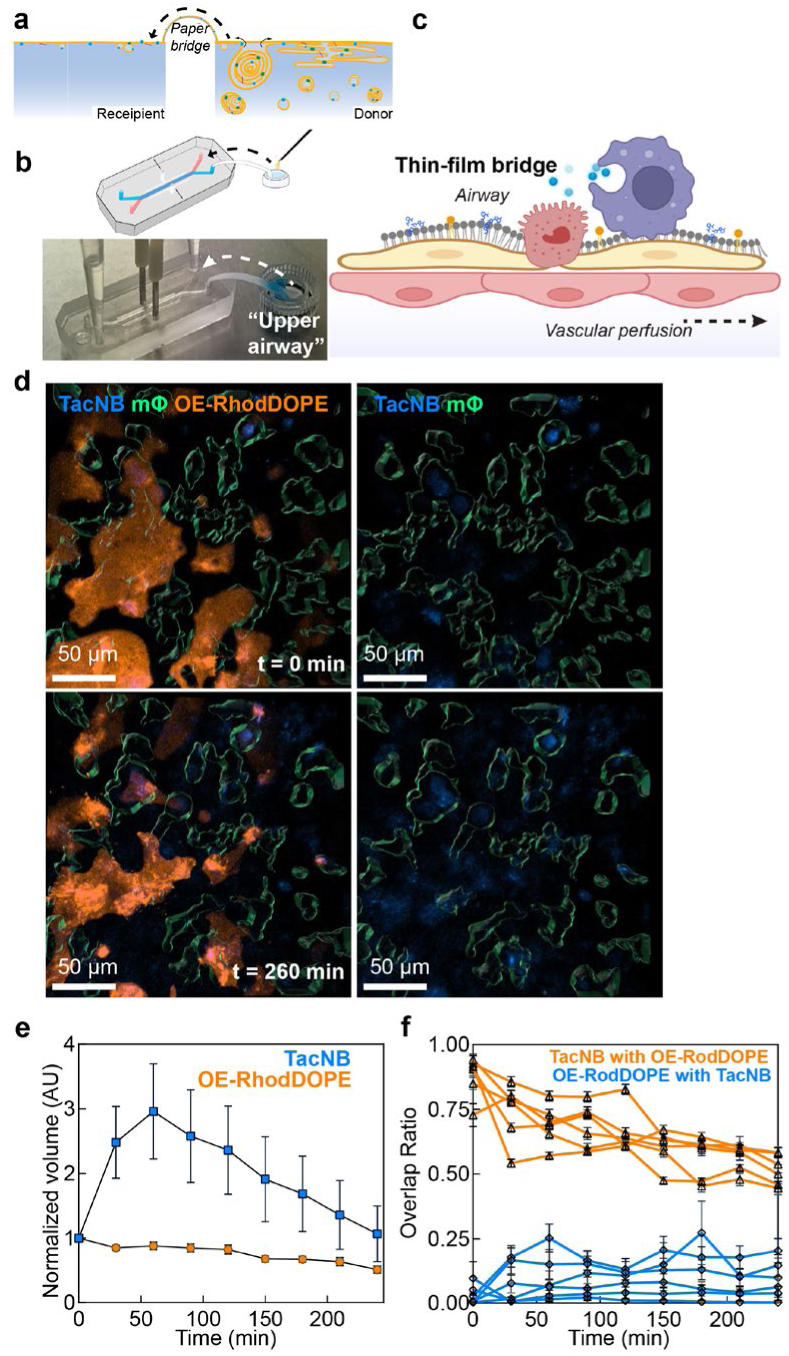
Recapitulation of vehiculation via thin-film bridge shows enhanced co-localization of cargo with pulmonary surfactant. (**a**) Schematic of vehiculation via thin-film bridge (TFB) in acellular models, adapted from ^43^. (**b**) Schematic and image of TFB vehiculation setup for the LoC model. (**c**) Schematic of the epithelial face of the LoC directly after TFB vehiculation. (**d**) Representative 3D views from live-cell confocal imaging of the epithelial face of a murine LoC. Left – OE-RhodDOPE (orange). Left and right – TacNB (blue), macrophages (green) are shown as surfaces. (**e**) Volume of TacNB and OE-RhodDOPE on the epithelial face normalized to the volume at t=0. Data from *n=6* fields of view. (**f**) Overlap ratio between TacNB and OE-RhodDOPE (orange) and the converse ratio between OE-RhodDOPE with TacNB (blue), data from *n=6* fields of view. Data shown is mean ± s.e.m.

Live-cell imaging in the murine LoCs on the microscope confirmed successful delivery of both surfactant lipids and the Tacrolimus cargo (Fig. 2d). Unlike for liquid administration, both TacNB and OE-RhodDOPE levels were relatively stable over the period of imaging (Fig. 2e). The volume of Tacrolimus was ca. 10-fold lower in the case of TFB vehiculation, but at later timepoints this was comparable across both delivery methods (Fig. S3). Levels of OE-RhodDOPE were comparable across the two delivery methods (Fig. S3). Both results are consistent with the performance of the TFB vehiculation in airway systems where surfactant lipids are vehiculated better than cargoes.

In a striking contrast with liquid administration, the TacNB co-localised extensively with the OE in both epithelial cells and macrophages (Fig.2f). TFB vehiculation increased the relative uptake in macrophages (Fig. S4) but there continued to be a relatively high volume taken up by epithelial cells.

### Alveolar macrophage-like cells in a human LoC recreate physiological delivery of functionally active Tacrolimus

We hypothesized that the impaired uptake of OE in the macrophages in the murine LoC was due to their inadequate recapitulation of the tissue-resident macrophage phenotype. Several recent studies have shown that circulating monocytes can differentiate into tissue-resident alveolar macrophages over time, driven by local cues ^24^. A recent protocol with human PBMCs has established alveolar macrophage like cells (AMLs) using a differentiation cocktail of GM-CSF, TGF-β, IL-10 and exposure to surfactant lipids to mimic the alveolar environment ^25^. We adopted a similar approach in this study, reasoning that media developed for the culture of human alveolar organoids ^26,27^ could be adapted into one for macrophage differentiation. Working exclusively with primary human cells from here on, we used a modified version of the alveolar organoid medium without the WNT activator CHIR99021 (K+DCI medium) further supplemented with 20 ng/mL of recombinant human GM-CSF ^28^, 10 ng/mL of recombinant human TGF-β ^29^, and 5% human serum. RNA-seq showed a much closer pattern of expression of genes characteristic of alveolar macrophages in these AMLs, relative to M-CSF differentiated macrophages (MDMs) (Fig. 3a). Unlike MDMs, the AMLs also showed enhanced uptake of exogenously provided pulmonary surfactant (Fig. 3b) and nuclear localisation of PPARγ ^30^(Fig. S5a). At 3 hours post-stimulation with lipopolysaccharide (LPS) from *Escherichia coli*, AMLs and MDMs showed upregulation of early NF-κB mediated genes *TNFA*, *IL1B*, and *IL6*, with a return to baseline at 24 hours post-stimulation (Fig. S5b), characteristic of TLR4 activation by LPS. In contrast, AMLs and MDMs differed significantly in response to infection with *Mycobacterium tuberculosis* (Mtb), the causative agent of tuberculosis (TB). AMLs showed higher infectivity (Fig. S5c) and permissivity for Mtb growth (Fig. S5d). At 4 dpi, AMLs but not MDMs showed increased expression of several genes including *SLC11A1*, *HMOX1*, and *PPARG* characteristic of the in vivo responses of alveolar macrophages in Mtb infection ^31,32^ (Fig. S5e).

**Fig. 3.**
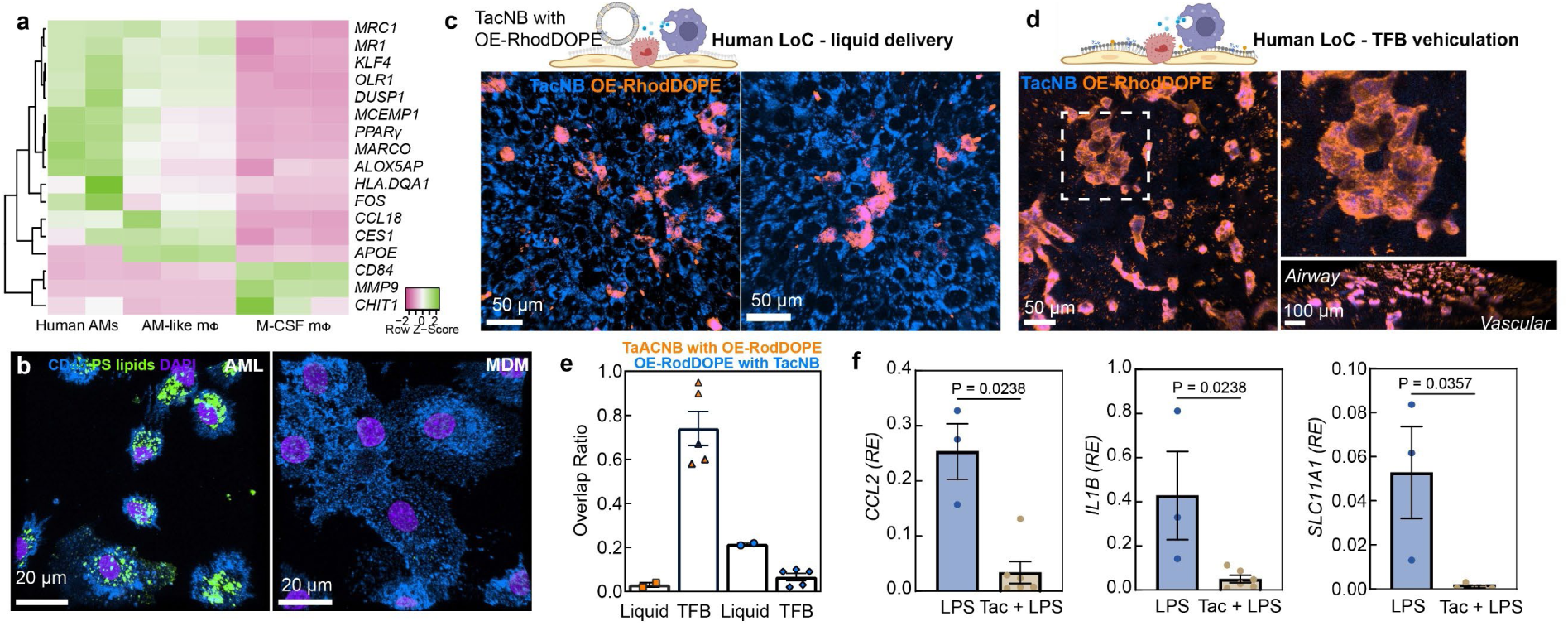
TFB vehiculation efficiently delivers surfactant lipids and functional cargoes to alveolar macrophages. (**a**) Heatmap of expression of selected markers of alveolar macrophages in alveolar-like macrophages (AMLs) and M-CSF differentiated macrophages (MDMs) from this study. Data for human AMs taken from ^25^. (**b**) Representative confocal images of AMLs and MDMs exposed to Curosurf^TM^ with 2% w/v RhodDOPE. (**c, d**) Representative confocal images of the epithelial face of human LoCs with TacNB (blue) delivered with OE-RhodDOPE (orange) in liquid formulations (**c**) or via thin-film bridge (TFB) delivery vehiculation (**d**). Zooms of the indicated area (white dashed square) and a perspective view are also shown. (**e**) Overlap ratio between TacNB and OE-RhodDOPE (orange) and between OE-RhodDOPE with TacNB (blue), data from *n=2* fields of view in liquid and n=6 field of view in TFB vehiculation. (**f**) Expression of indicated genes relative to *GAPDH* measured by qRT-PCR of cells in the epithelial channel of human LoCs exposed to LPS, or with Tacrolimus delivered via TFB vehiculation prior to LPS exposure. Data from *n=3* control LoCs and *n=5* TFB-vehiculated LoCs, respectively. Data shown is mean ± s.e.m. P-values calculated by a two-tailed Mann-Whitney test (**f**).

We therefore tested the delivery of TacNB to human LoCs, reconstituted as described previously ^33,34^, but with the addition of AMLs in place of MDMs. Even in this model, the administration of OE with TacNB in liquid resulted in uptake overwhelmingly in the alveolar epithelial cell layer, with little or no uptake in alveolar macrophages (Fig. 3c) and poor co-localisation between OE and TacNB (Fig. 3e). In contrast, TFB-vehiculation showed TacNB localized overwhelmingly in the AMLs, identified clearly by their more amoeboid cell shape on top of the underlying epithelial layer (Fig. 3d, inset). Within these cells, TacNB co-localised almost exclusively with Rhod-DOPE. Very little TacNB was observed in underlying alveolar epithelial cells. In contrast, smaller volumes of OE-RhodDOPE were visible in the epithelial cells (Fig. 3e). Thus, the airway delivery via thin-film bridge in human LoCs reconstituted with AMLs appeared closest to recapitulating the in vivo delivery of exogenous substances via surfactant vehiculation, demonstrating significant uptake in macrophages.

Lastly, we examined, if Tacrolimus delivered via TFB vehiculation at the air-liquid interface was functionally effective, by evaluating efficacy in reducing inflammatory responses to sterile inflammation. We used the unconjugated Tacrolimus formulation to ensure effective immunological function. First, the successful delivery of surfactant lipids was verified by confocal imaging. 30 minutes later, the epithelial face of the LoC was exposed to LPS for 1 hour. In untreated controls, the epithelial face of the LoC was directly exposed to LPS. Transcriptional profiling via qRT-PCR of the total RNA from cells from the epithelial channel 3 hours after the removal of LPS solution showed a marked reduction in inflammation in LoCs with pre-exposure to Tacrolimus (Fig. 3f).

### TFB vehiculation shows proof-of-concept for functional efficacy of aerosolized TB antibiotics

The airway delivery of antibiotics represents an untapped route for pulmonary infections, particularly chronic ones caused by non-tuberculosis mycobacteria such as *Mycobacterium abscessus* ^35, 36^ or TB^37^. Several antibiotics effective against mycobacteria are lipophilic – this makes delivery via a vascular route to pulmonary sites of infection less efficient, but is an ideal characteristic for aerosolized therapy^38^. A significant limitation for further development and optimization of aerosolized therapy is the unavailability of effective in vitro models of the delivery and uptake. We therefore designed a set of proof-of-concept experiments applying thin-film bridge approach to demonstrate vehiculation of bedaquiline (BDQ), a frontline antibiotic against *Mycobacteria spp* ^39–41^. In these formulations, the OE was labelled with Cy5-PC (Fig. 4a), but the BDQ was unlabelled. This change in cargo did not appear to alter the efficacy of vehiculation. Assuming similar dynamics as for TacNB, we hypothesized that sites with OE-Cy5PC accumulation also represented cells that had taken up BDQ and were overwhelmingly AMLs. Three hours later, the LoCs were infected with Mtb Erdman constitutively expressing tdTomato, as described earlier ^33^. Bacterial microcolonies were then imaged 1 and 2 days-post infection with live-cell widefield microscopy, and growth imputed from changes in microcolony size (Fig. 4b). Zooms in Fig. 4c show an area of control of bacterial growth which co-localises with OE-Cy5PC signal, whereas the zooms in Fig. 4d show unrestrained bacterial growth in cells without OE-Cy5PC signal. Overall, bacterial microcolonies in cells that were OE-Cy5PC+ showed consistently lower growth rates than bystander cells which did not take up the exogenously provided formulation, where growth rates were consistent with untreated control LoCs (Fig. 4e). This data is the first demonstration that BDQ can be successfully vehiculised with surfactant formulations, that BDQ colocalizes with surfactant formulations in alveolar cells (likely accumulating in macrophages that are themselves the cells that take up Mtb in the lung) and that the delivered antibiotic is effective even 48 hours after administration.

**Fig. 4.**
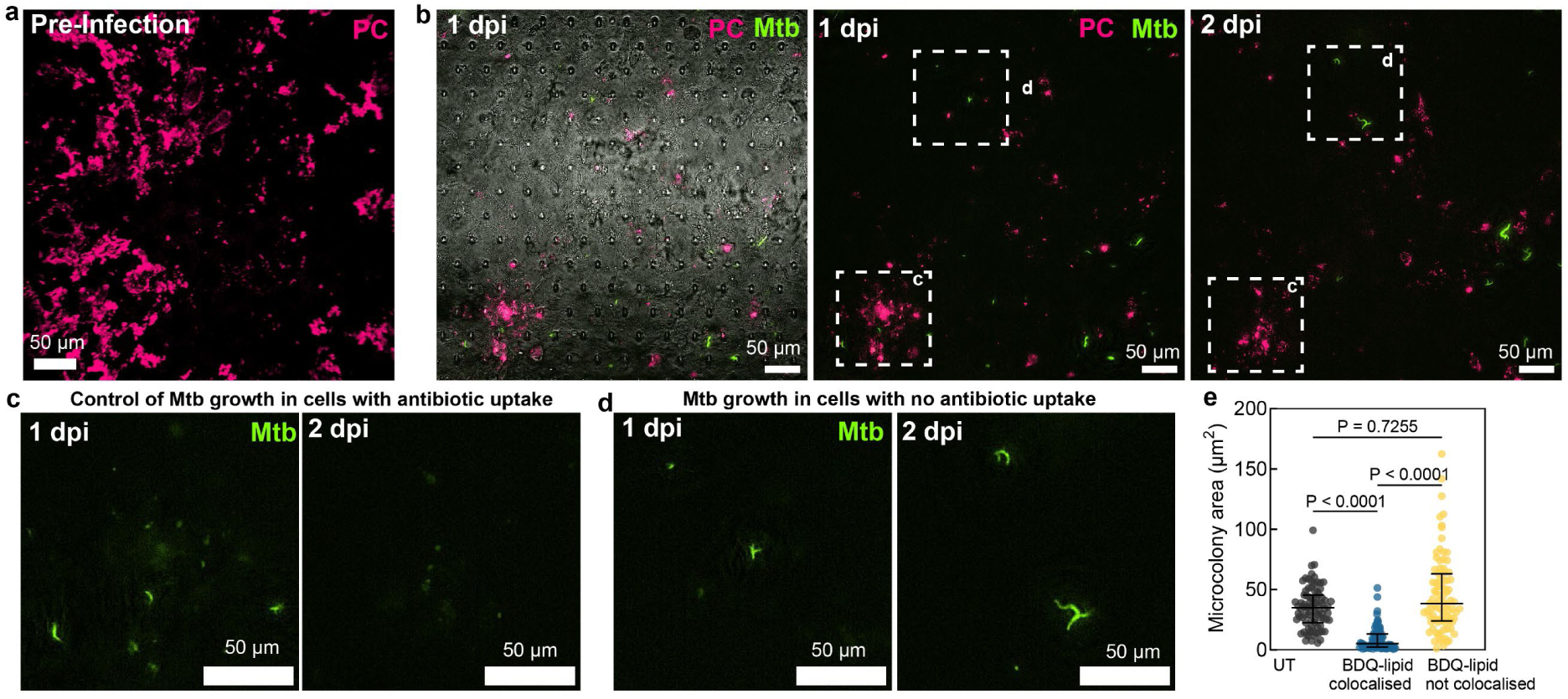
Prophylactic delivery of Bedaquiline via TFB vehiculation attenuates Mtb growth. **a,** Confocal image of the epithelial face of a human LoC after TFB-vehiculation of bedaquiline (BDQ) with OE-Cy5PC (pink). (**b**) Extended depth of focus snapshots from time-lapse imaging of human LoCs at 1 and 2 days post-infection (dpi) with Mtb constitutively expressing tdTomato (yellow). (**c**, **d**) Zooms from the indicated regions in (**b**), only the Mtb channel is shown. (**e**) Quantification of Mtb growth in untreated LoCs (*n=2* LoCs), and areas on LoCs with prior BDQ vehiculation where Mtb and BDQ-OE-Cy5PC co-localized or did not (*n= 3* LoCs). Data shown is mean ± s.e.m. P-values calculated by a one-way Kruskal-Wallis ANOVA with Dunn’s multiple comparison test (**e**).

## Discussion

The alveolar interface is a highly dynamic environment - populated with a defined small number of cell types, constantly exposed to mechanical stimulation in addition to a unique tissue barrier with the ambient air. This barrier must be as thin as possible to allow gas exchange. The unique properties of this barrier are made possible by the alveolar lining fluid and the layer of pulmonary surfactant at the interface, whose biophysical properties are closely intertwined with its biochemical characterization. In the past several years, air-liquid interface culture of epithelial cells, including in lung-on-chip models has become *de rigeur*. This approach has provided several new insights into epithelial cell biology and alveolar immune responses at the air-liquid interface and driven advances in cell culture techniques including co-culture to enable better recapitulation of cell types. However, the vast potential for these systems to explore the unique biophysical nature of the alveolar lining fluid at the air-liquid interface remains underutilized.

In this work, we have systematically compared delivery strategies in vitro, using small lipophilic compounds as cargo. Even in unphysiological liquid inoculations, incorporation of the cargo with a pulmonary surfactant extract resulted in a substantially enhanced uptake. This could be because of receptor-mediated uptake of specific lipid components of surfactant, as enhanced uptake was also observed for multi-lamellar suspensions lacking proteins SP-B or SP-C. Further work is needed to understand cell-type specific uptake mechanisms, and how the lipophilic molecules rapidly separate from surfactant components once internalised within the cell. In contrast to this, we develop and demonstrate a feasible mechanism for mimicking vehiculation of surfactant lipids and cargo over a thin-film bridge, an approach that has only been used so far in acellular experiments. Notably, the stretching dynamics that mimic breathing were essential to drive the vehiculation process, similarly to what it was already shown in interfacially-interconnected acellular compartments ^10^, interpreted as the importance for interface-assisted vehiculation of surface-tension driven Marangoni flows. Although this transported less cargo overall than exposure to liquid formulations, not only did the cargo remain associated with the surfactant lipids, but was also retained over a longer period at the airways.

In further methodological innovation, we showed that macrophages differentiated in vitro to more closely resemble alveolar macrophages performed the function of surfactant uptake to a greater degree than those we have used previously. In human LoCs reconstituted with alveolar-like macrophages, but only for TFB vehiculation, we found that most of the surfactant lipids and cargo were delivered to and persisted in these cells. Our systematic study therefore highlights the importance of recreating both the biophysics and the cell phenotypes for models of alveolar delivery. The long-term retention of delivered cargoes is an important observation in our study. We directly demonstrate that through this, the delivered cargoes have prophylactic activity. In the case of Tacrolimus, this blunted inflammatory responses to subsequent LPS exposure. Extending this to the lipophilic antibiotic bedaquiline, we could show that a one-time delivery had an antimicrobial effect even up to 48 hours later. Both TB and NTM infections suffer from a severe pill burden and associated toxicities from multi-drug regimens over several months. Our proof-of-concept experiments suggest that vehiculation using pulmonary surfactant could deliver long-acting antimicrobial formulations in a localised manner to the pulmonary site of infection.

Although this study has focused on small molecule cargoes, it is likely that similar dynamics govern the vehiculation of particulate matter including viruses and bacteria delivered by aerosols. In addition, the filter paper bridge favours the vehiculation of the surfactant layer but hinders the movement of the surface-associated reservoirs located just below the interface^42^. Furthermore, this mechanism likely misses biological cues delivered by the underlying epithelium during the transport process. Future studies will develop approaches to recapitulate this to a greater extent,

## Acknowledgements

The authors wish to thank members of the BioImaging and Optics Platform (BIOP) Core Facility, the Bioinformatics Core Facility and Prof John D McKinney at EPFL, Annette Fautsch for experimental assistance and Prof Guilllermo Orellana Moraleda for generation of fluorescently conjugated Tacrolimus and Beclomethasone. CG-M acknowledges scholarships from the Spanish Ministry of Science, Innovation and Universities (FPU16/02553 and EST19/00993). JP-G acknowledges support from the Spanish Ministry of Science and Innovation (PID2024-156556OB-I00). VVT acknowledges support from the Holcim Stiftung zur Förderung der Wissenschaftlichen, intramural support from the Medical Faculty of Heidelberg University and a Project Grant from the Stiftung zur Förderung der Erforschung von Ersatz- und Ergänzungsmethoden zur Einschränkung von Tierversuchen (SET).

## Author contributions

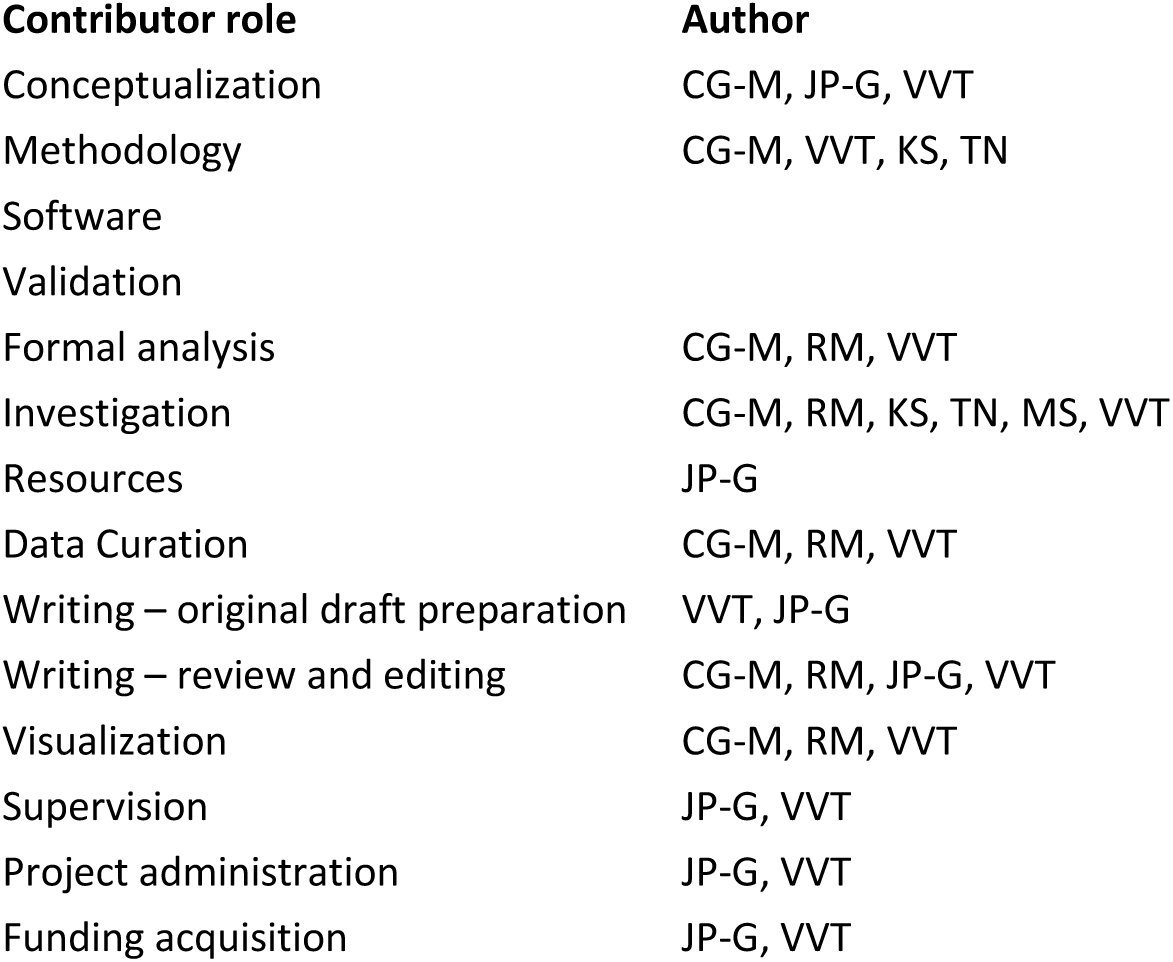

---

**Fig. S1.**
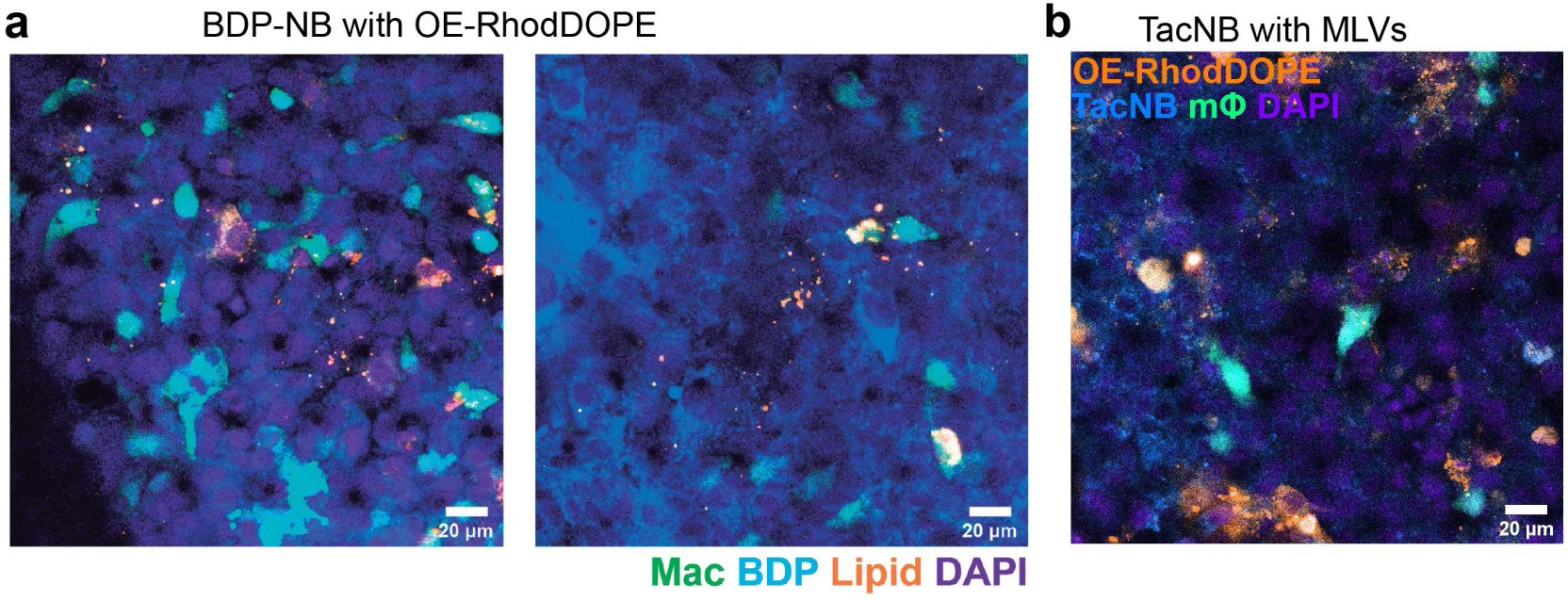
Liquid formulations with pulmonary surfactant enhances cargo uptake. **(a)**, Representative confocal images of the epithelial face of a murine LoC after delivery of beclomethasone dipropionate conjugated with Nile Blue (BDP-NB, blue) with OE-RhodDOPE (orange) by addition of liquid to the epithelial channel. (**b**) Representative confocal image of the epithelial face of murine LoC after delivery of TacNB (blue) with pulmonary surfactant multi-lamellar vesicles (MLVs, orange). Macrophages – green, DAPI – purple.

**Fig. S2.**
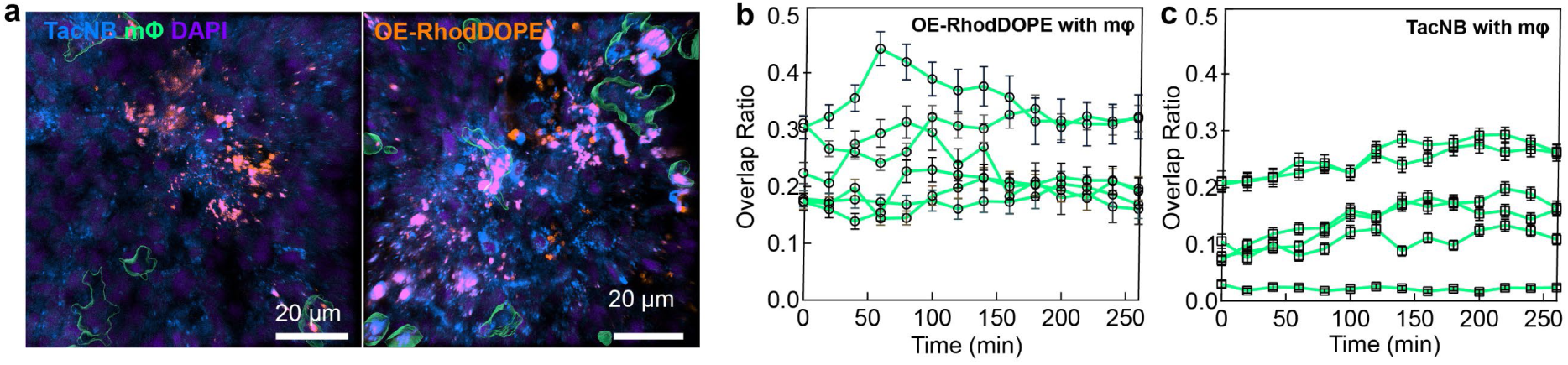
Additional characterization of delivery of Tacrolimus in liquid formulation with pulmonary surfactant. **(a)**, Representative confocal images of the epithelial face of a murine LoC after inoculation with TacNB with OE-RhodDOPE. TacNB – blue, macrophages – green, OE-RhodDOPE – orange. (**b, c**) Overlap ratio between TacNB (**b**) and OE-RhodDOPE (**c**) respectively with macrophages, data from *n=6* fields of view.

**Fig. S3.**
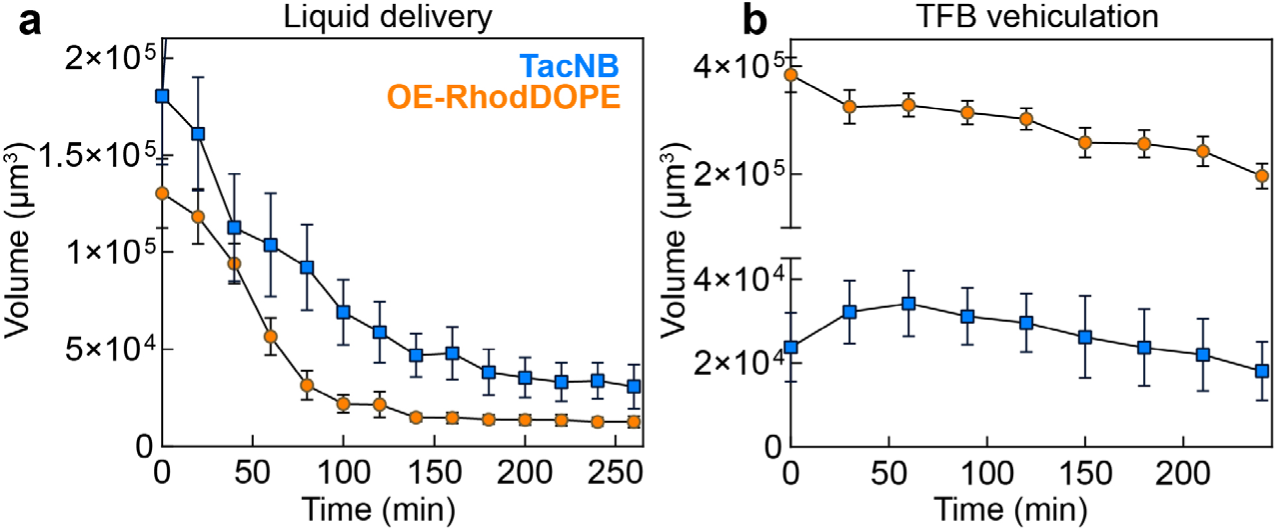
Additional characterization of delivery of Tacrolimus in liquid formulation with pulmonary surfactant vs TFB vehiculation. Volume of TacNB and OE-RhodDOPE on the epithelial face as a function of time in (**a**) liquid delivery, related to Fig. 1f and (**b**) TFB vehiculation, related to Fig. 2e. Data from *n=6* fields of view.

**Fig. S4.**
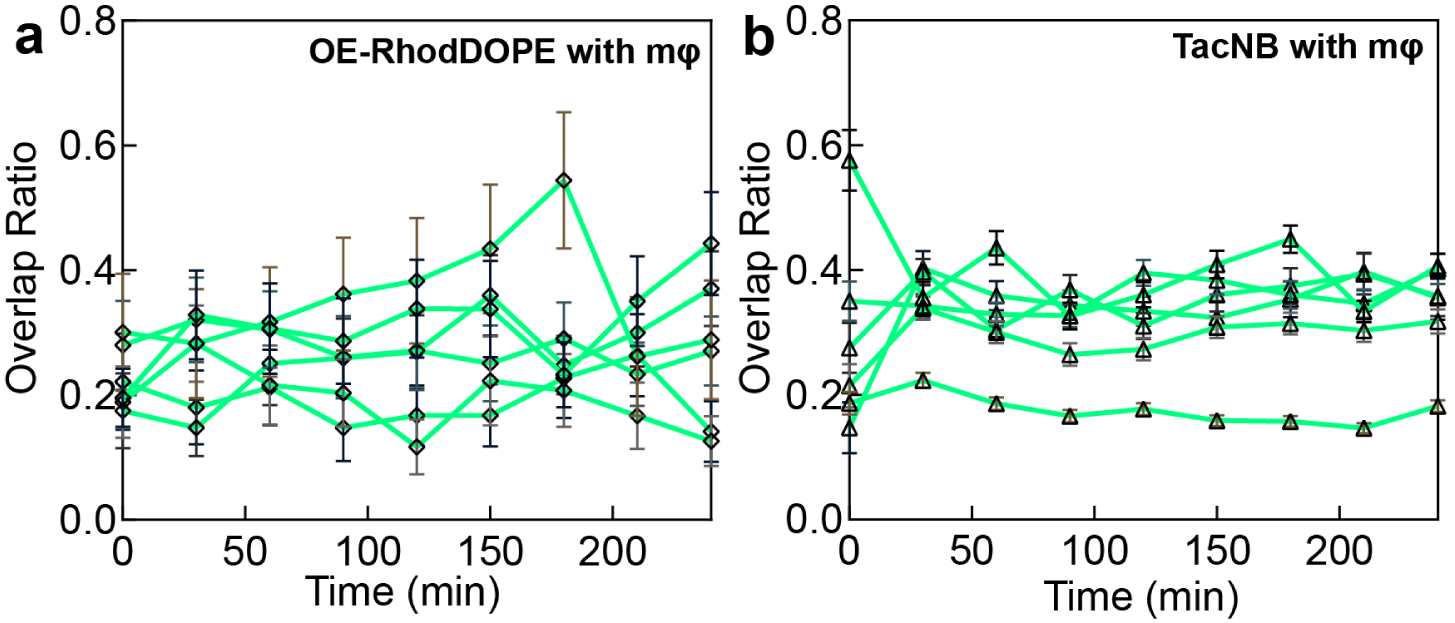
Additional characterization of delivery of Tacrolimus in TFB vehiculation. (**a, b**) Overlap ratio between TacNB (**a**) and OE-RhodDOPE (**b**) respectively with macrophages, data from *n=6* fields of view.

**Fig. S5.**
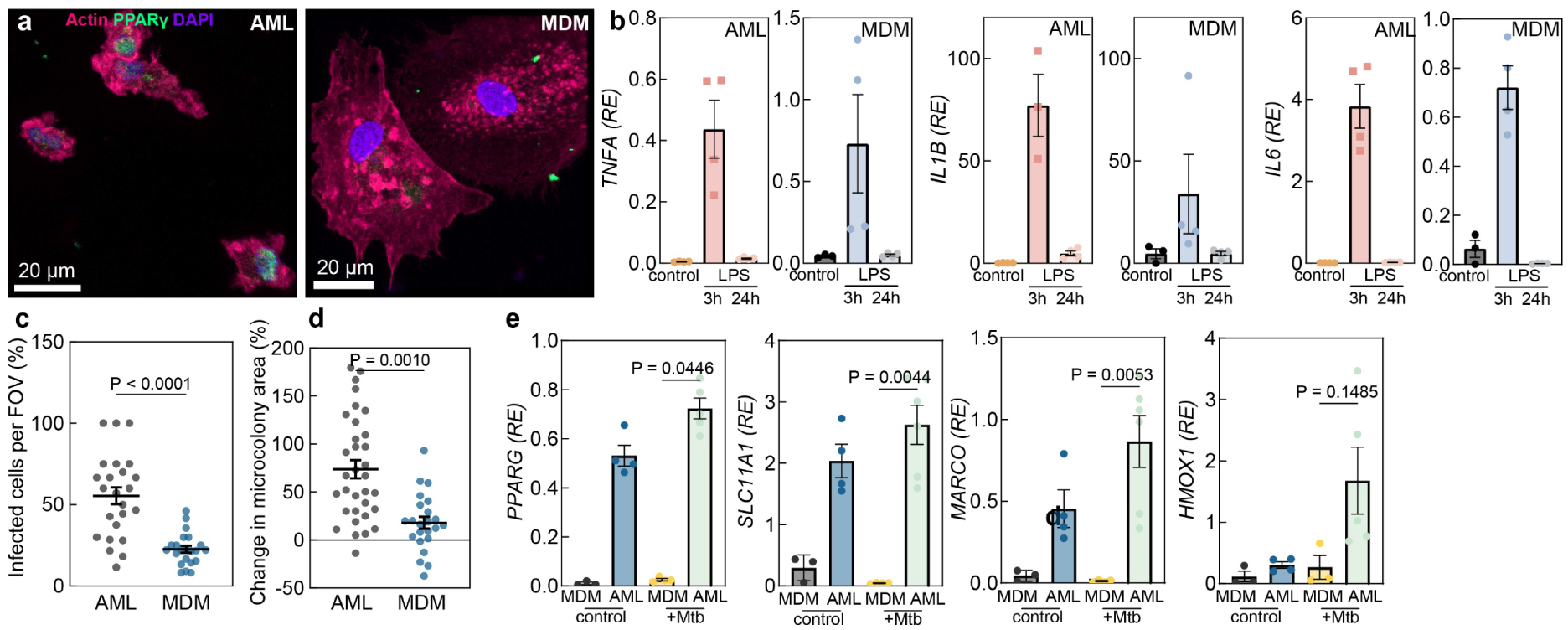
Additional characterization of delivery of Tacrolimus in TFB vehiculation. (**a**) Representative confocal images of AMLs and MDMs immunostained for PPARγ (green). Actin (pink), DAPI (purple). (**b**) Expression in AMLs and MDMs of indicated genes relative to *GAPDH* measured by qRT-PCR of unstimulated controls, and cells at 3 hours and 24 hours post-exposure to 100 ng/mL LPS. (**c**) Quantification of infection efficacy of AMLs and MDMs with Mtb. (**d**) Mtb growth rates characterized by live-cell imaging of microcolonies in AMLs and MDMs at four days post-infection. (**e**) Expression in AMLs and MDMs of indicated genes relative to *GAPDH* measured by qRT-PCR of uninfected controls, and cells infected with Mtb at 4 days post infection. Data shown is mean ± s.e.m. P-values calculated by a two-tailed Mann-Whitney test (**c, d, e**).

## Supplementary Legends

**Supplementary Table 1:** List of primers for qRT-PCR used in this study.

**Supplementary Table 2:** List of antibodies used in this study.

**Supplementary Note 1:** Surfactant formulations used in this study.

## Materials and Methods

### Alveolar epithelial cell and lung microvascular endothelial cell culture

Primary murine and human alveolar epithelial cells (ATs) and murine and human lung microvascular endothelial cells were obtained from Cell Biologics, USA. All cell types were cultured in vitro in the appropriate complete medium comprising base medium and supplements (Cell Biologics, USA) in 5% CO_2_ at 37 °C as previously described ^33,34^. All lung-chips were reconstituted with ATs seeded directly on the LoC, without any additional *in vitro* culture. Endothelial cells were passaged between 3-5 times before seeding in the LoC devices. In the case of human LoCs, experiments were performed with cells from at least two donors.

### Murine macrophage culture

For murine bone marrow-derived macrophages (BMDMs), bone marrow was obtained from the femurs of 6- to 8-week-old female C57BL/6 mice (Charles River) or the C57BL/6-Tg(CAG-EGFP)131Osb/LeySopJ (Jackson Labs) mice that constitutively express GFP in every cell and cryopreserved. Animal protocols were reviewed and approved by EPFL’s Chief Veterinarian, by the Service de la Consommation et des Affaires Vétérinaires of the Canton of Vaud, and by the Swiss Office Vétérinaire Fédéral under license VD 3434. Bone marrow was cultured in Dulbecco’s Modified Eagle Medium (DMEM) (Gibco) supplemented with 10% FBS and 20% L929-cell-conditioned medium (as a source for macrophage-colony stimulating factor protein or M-CSF) for 7 days. No antibiotics were used in the cell culture media for all cell types to avoid activation of macrophages or inhibition of Mtb growth. Thereafter the differentiated macrophages were washed twice with PBS, and detached with a cold solution of 2 mM EDTA in PBS. Detached cells were recovered in murine alveolar epithelial media without dexamethasone and introduced onto the epithelial face of the LoCs.

### Human macrophage culture

Peripheral blood mononuclear cells (PBMCs) were obtained from buffy coat (Interregional Blood Transfusion SRC Ltd, Switzerland, or Institut fuer Klinische Transfusionsmedezin und Zelltherapie (IKTZ), Heidelberg) taken from anonymized donors. PBMCs were isolated using a Biocol Separation procedure as per the manufacturer’s instructions. Isolated PBMCs were subsequently cryopreserved in a solution of 70% heat-inactivated fetal bovine serum (FBS, Gibco), 20% RPMI medium (Gibco) and 10% dimethyl sulfoxide (DMSO). One week prior to seeding the macrophages in the LoC devices, a cryopreserved aliquot was thawed and cultured in a T-75 flask (TPP, Switzerland) in RPMI supplemented with 10% FBS. CD14+ monocytes were isolated using positive selection (CD14 Ultrapure Isolation Kit, Miltenyi Biosciences), embedded in hemispherical domes of basement membrane extract (BME-2, Cultrex) in 24 well plates.

#### MDM differentiation

CD14+ monocytes in hemispherical BME-2 domes were cultured in DMEM medium supplemented with 10% FBS and differentiated for 7 days with 20 ng/mL recombinant human macrophage-colony stimulating factor protein (M-CSF) (Thermo Fisher Scientific).

#### AML differentiation

CD14+ monocytes in hemispherical BME-2 domes were cultured in a modified version of the cell culture medium used in Burgess et al.^44^ The medium consists of complete serum-free medium (cSFDM) with a reduced amount of Primocin (at 200 ng/mL) supplemented with 10 ng/mL recombinant human Keratinocyte Growth Factor (KGF), 50 nM dexamethasone, 0.1 mM 8-Bromoadenosine 3’,5’-cyclic monophosphate (8-BrcAMP), and 0.1 mM 3-isobutyl-1-methylxanthine (IBMX) to make the K+DCI media. This formulation is then supplemented additionally with 5% autologous serum (or serum from AB donors), 20 ng/mL human granulocyte-macrophage colony-stimulating factor (GM-CSF) and 10 ng/mL human transforming growth factor-beta (TGF-β).

#### Removal of differentiated macrophages

Differentiated macrophages were isolated by disrupting the BME-2 domes on ice. The BME-RPMI suspension was then centrifuged at 200 g for 5 minutes in a 15 mL tube pre-coated with 1% BSA in PBS, resuspended in 4-5 mL of ice-cold Cell Recovery Solution (Corning) per well, and incubated at 4°C for 30 minutes. The cell suspension was then washed with 10 mL RPMI medium supplemented with 10% FBS to remove any remaining traces of the recovery reagent. An additional digestion step with cold Cell Recovery solution (Corning) was then performed until no visible matrix fragments were obtained when the cell solution was centrifuged at 300g. The cell suspension was then washed with 10 mL RPMI medium supplemented with 10% FBS to remove any remaining traces of the recovery reagent. Isolated macrophages were resuspended in a small volume of epithelial cell media without dexamethasone and introduced onto the epithelial face of the LoCs.

### RNAseq of differentiated macrophages

Human MDMs and AMLs were differentiated in BME-2 domes as described above. At the end of the differentiation period, the media was aspirated and the domes were homogenized in RLT plus buffer from the Qiagen RNeasy Plus Micro kit supplemented with β-mercaptoethanol, and RNA isolated as described below.

Libraries were prepared from ribodepleted RNA isolated from macrophages and sequenced using the NEBNext^®^ Ultra^™^ II RNA Library Prep Kit for Illumina^®^ (New England Biolabs) as per the manufacturer’s instructions. A mean of 77.7 million sixty base-pairs paired-end reads were obtained among the six samples (72.1 – 86.0 million reads). The quality of sequenced Fastq files were analyzed using FastQC (version 0.11.9) and reads were mapped on the reference Human genome using STAR (v2.7.10b) ^45^. The uniquely mapped reads were between 80.3%-83.9%. Reads count per gene was performed by STAR using option *--quantMode GeneCounts*.

Gene expression quantification was performed using R package edgeR ^46^. Genes below a mean of ten reads count among the samples were filtered out. Libraries normalization was performed using the TMM method and gene expression quantified as count per million (CPM).

### Immunofluorescence staining of MDMs and AMLs

For immunofluorescence labelling, AMLs or MDMs were fixed by immersing completely in a freshly diluted solution of 4% (PFA) in PBS for 2 hours at room temperature followed by two washes with PBS. The fixed sample was permeabilized with 0.1% Triton X-100 (Sigma-Aldrich) and 2% saponin (Sigma-Aldrich) in PBS for 15 minutes at room temperature followed by incubation with a blocking solution of 2% bovine serum albumin (BSA) in PBS (“blocking buffer”). The permeabilized and blocked sample was then incubated overnight with the primary antibody in the blocking buffer at 4°C. A list of primary and secondary antibodies and concentrations used is included in Table S2. The following day, the sample was washed extensively with fresh blocking buffer before incubation with the secondary antibody at a dilution of 1:300 for 1 hour at room temperature. Infected macrophages or LoCs were also stained with the Hoechst dye (Thermo Fisher) at 1:1000 dilution for 30 minutes at room temperature for visualizing nuclei. Confocal images were acquired on an inverted Leica SP8 microscope.

### Murine and human LoC models

The protocols for both murine ^47^ and human ^48^ LoCs have been previously reported and were implemented as describe in this study. Once the epithelial and endothelial cell layers were matured in co-culture, macrophages were introduced onto the epithelial face. The LoC was incubated for 30 minutes (for BMDMs in murine LoCs) or overnight (for AMLs in human LoCs) at 37°C and 5% CO_2_ to allow macrophages to attach to the ATs. Thereafter the medium on the epithelial face was then removed, and the chip was maintained overnight at an air-liquid interface (ALI). Chips were controlled to ensure that they successfully maintained the ALI overnight and then used for further experiments. No antibiotics were used in any of the cell culture or flow media for infection experiments.

### Generation of pulmonary surfactant organic extract formulations

Organic extracts (OE) of pulmonary surfactant purified from porcine bronchoalveolar lavages were prepared as previously described ^49,50^. DPPC, POPG, Rhodamine labelled DOPE (RhodDOPE), Cy5-labelled PC (Cy5PC), Tacrolimus, and Bedaquiline (BDQ) are obtained commercially. Tacrolimus conjugated to Nile Blue and Beclomethasone conjugated to Nile Blue were generated as explained elsewhere ^10,42^. Appropriate volumes of OE, Rhodamine labelled DOPE or Cy5 labelled PC, Tacrolimus, Tacrolimus conjugated to Nile Blue, Beclomethasone conjugated to Nile Blue or bedaquiline (see Supplementary Note 1) were dried in a 1.5 mL Eppendorf tube under a nitrogen stream. Once dried, the nitrogen flux was increased to ensure complete evaporation of the organic solvent. Next, an aqueous suspension of surfactant-containing solutions was generated by hydrating the lipid film with an appropriate volume of PBS. The suspension was incubated at 45 °C in a thermomixer for 1 hour with shaking every 10 minutes. This temperature is above the melting temperature of surfactant phospholipids as well as that for the multilamellar vesicles made of 7:3 DPPC: POPG. Thereafter the samples were stored at 4 °C for a short period before being used for experiments within the subsequent 24 hours.

### Liquid delivery of pulmonary surfactant organic extract containing formulations in murine and human LoCs

For delivery via liquid, the samples prepared at 5 mg/mL of phospholipids in PBS were further diluted in the murine or human epithelial medium to a concentration of 2 mg/mL. 30 µL was added to the epithelial channel and the LoCs incubated at 37°C for 30 minutes without vascular perfusion. Thereafter, the inoculum was removed and the LoC returned to air-liquid interface and vascular perfusion. For TacNB only delivery, TacNB was dried under nitrogen as described above, dissolved in DMSO at 2.5 µg/µL, and then diluted 25-fold in PBS. The solution in PBS was then added to epithelial cell medium in the same manner as described above.

### Thin-film bridge delivery of pulmonary surfactant organic extract containing formulations in murine and human LoCs

Pulmonary surfactant organic extract containing formulations were generated as described above. Small wedges of Whatman filter paper were generated, and sterilized for 20 min under the UV before use. The plastic cap of a 1.5 mL O-ring tube (Sarstedt) was used as a donor trough. The narrow-end of the wedge-shaped filter was placed into the epithelial channel of the LoC with tweezers. The remaining portion of the wedge was then hydrated carefully with ca. 100 – 200 µL until the broad end of the wedge contacts the surface of the liquid in the donor trough. Thereafter, any excess liquid from the donor trough was removed to retain ca. 150 µL volume. To this donor trough, 10 µL of the sample was carefully added at the interface to minimise addition to the bulk of the liquid wherever possible. The LoC was maintained under breathing conditions but without perfusion using a microfluidic pressure regulator (Elveflow, OB1 Mk4) for the duration of the bridge administration (30 minutes). Thereafter the LoC was returned to regular vascular flow conditions.

### Confocal imaging of LoCs to study delivery of surfactant containing formulations

LoCs were placed in a microscope stage-top incubator and mounted on the stage of the Leica SP8 or Leica Stellaris 5 confocal microscopes. The stage-top incubator was connected to a gas mixer (Okolab) to maintain 5% CO_2_ and a temperature of 37°C throughout the imaging period. For LoCs, flow of medium through the vascular channel was maintained throughout this period via the use of a syringe pump. The LoC was imaged using a 25x water immersion objective (NA = 0.95, Leica) for snapshot imaging. The epithelial face of the LoC (where the refractive index differences is highest due to the ALI) was maintained in focus. At each timepoint, a Z-stack with an axial spacing of 0.3 μm was taken for a series of fields of view along the entire length of the LoC. Bright-field images were also captured.

### Confocal image analysis

Z-stacks were subsequently deconvolved using the Huygens Deconvolution Software (Scientific Volume Imaging). Imaris 9.9 (Bitplane) was used to analyse the intensity of immunofluorescence. Imaris was also used for rendering 3D images.

### Mycobacterium tuberculosis culture

The Mtb Erdman strain constitutively expressing the fluorescent protein tdTomato ^20,33^ was cultured at 37°C in Middlebrook 7H9 (Difco) supplemented with 0.5% albumin, 0.2% glucose, 0.085% NaCl, 0.5% glycerol, and 0.02% tyloxapol for liquid culture and Middlebrook 7H11 (Difco) supplemented with 10% OADC enrichment (BD) and 0.5% glycerol for solid culture. Aliquots were stored in 15% glycerol at −80 °C and used once.

### Evaluation of transcriptional responses of human MDMs and AMLs to LPS

Human MDMs and AMLs were differentiated in BME-2 domes as described above. Isolated macrophages were seeded overnight in 24 well plates (Corning) in their corresponding differentiation media. The next day, the media was refreshed and the macrophages stimulated with LPS for 3 hours or 24 hours. At the desired timepoint, the media was aspirated and RNA collected using reagents from the Qiagen RNeasy Plus Micro kit as described to evaluate transcriptional responses via qRT-PCR.

### Evaluation of transcriptional responses of human LoCs to LPS

Human LoCs were reconstituted with AMLs as described above and were provided Tacrolimus delivered via TFB vehiculation for 30 minutes. Thereafter the LoCs were returned to perfusion for 30 minutes at 37°C, 5% CO_2_. At this point, a solution of LPS at 100 ng/mL in human alveolar epithelial cell media without FBS was introduced into the epithelial channel and the LoC incubated at 37 °C, 5% CO_2_ without vascular perfusion. After 60 minutes, the media in the epithelial cell channel was withdrawn and the LoC returned to air-liquid interface, and maintained under perfusion at 3 7°C, 5% CO_2_ for a further three hours. At this point, RNA was extracted separately from the epithelial channel and the endothelial channel as previously described ^20,33,34^ and analysed further via qRT-PCR. For untreated controls, the steps prior to LPS stimulation were omitted.

### Infection of human AMLs with Mtb

Human AMLs were seeded overnight in 24 well plates (Corning) in AML media. The next day, a 1 mL aliquot of an Mtb-tdTomato culture grown to exponential phase (OD_600_ 0.3-0.5) was centrifuged at 5000 g for 5 minutes at room temperature, the supernatant was removed, and the cell pellet was resuspended in 200 μL AML media. A single-cell bacterial suspension was generated via filtration through a 5-μm syringe filter (Millipore). The single-cell suspension was diluted 100-fold in AML media and 100 μL added to the wells of the well plate. The cells were incubated for 3 hours at 37 °C and 5% CO_2_ to allow Mtb infection of cells after which the inoculum was withdrawn, the macrophages washed with PBS and then incubated with fresh AML media. The media was refreshed every 2 days. RNA was isolated at 4 days post infection.

### Infection of human LoCs with Mtb

LoCs either pre-exposed to formulations of bedaquiline containing pulmonary surfactant delivered via TFB vehiculation or untreated controls were assembled into a stage top incubator (Okolab) prior to infection and flow of medium through the vascular channel was maintained throughout the course of the experiment by use of a syringe pump. A 1 mL aliquot of an Mtb culture grown to exponential phase (OD_600_ 0.3-0.5) was centrifuged at 5000 g for 5 minutes at room temperature, the supernatant was removed, and the cell pellet was resuspended in 200 μL of epithelial cell media without FBS. A single cell suspension was generated via filtration through a 5-μm syringe filter (Millipore). The single-cell suspension was diluted 100-fold in epithelial media and 30 μL was added to the epithelial channel of the LoC. chip was incubated for 2-3 hours at 37 °C and 5% CO_2_ to allow Mtb infection of cells on the epithelial face, after which the solution on the epithelial face was withdrawn. The epithelial face was returned to ALI, and the inlets of the infected chip were sealed with solid pins as a safety precaution for incubation and imaging in the BSL-3 facility.

### Widefield imaging of human AMLs and human LoCs to measure Mtb growth

LoCs or infected AMLs in 24-well plates (Ibidi) were placed in a microscope stage-top incubator and mounted on the stage of the Leica Thunder widefield microscope. The stage-top incubator was connected to a gas mixer (Okolab) to maintain 5% CO_2_ and a temperature of 37 °C throughout the imaging period. For LoCs, flow of medium through the vascular channel was maintained throughout this period via the use of a syringe pump. The LoC was imaged using a 25x water immersion objective (NA = 0.95, Leica) for snapshot imaging. The epithelial face of the LoC (where the refractive index differences is highest due to the ALI) was maintained in focus. At 1 and 2 days-post-infection, a Z-stack with an axial spacing of 2.5 μm was taken for a series of fields of view along the entire length of the LoC. Bright-field images were also captured. For AMLs, higher resolution imaged with a 60x oil immersion objective (NA=1.4) were obtained immediately after infection (to evaluate infectivity) and then for the subsequent 4 days post infection to evaluate growth rates.

### Widefield imaging and image analysis

Z-stacks were processed using the Lightning deconvolution capability of the Leica Thunder microscope. ImageJ was used to analyse the size of Mtb microcolonies, and to render extended depth of focus images for display.

### RNA isolation and quantitative real-time PCR (qRT-PCR)

For RNA isolation from Mtb-infected macrophages in well plates, 800 μL of TRIzol Reagent was added to samples per well and collected by mechanical scraping. The cell lysate was mixed with 200 μL chloroform and centrifuged at 12000 g for 15 minutes at 4°C. The top aqueous phase was collected and mixed with an equal volume of isopropanol and brought out of the BSL-3 for RNA purification.

For RNA isolation from eukaryotic cells in LoCs samples were collected from apical and vascular channels separately in approximately 350 μL of the RLT Plus buffer of the Qiagen Micro Plus RNA Isolation Kit and RNA was purified as per the manufacturer’s instructions.

cDNA was generated using the SuperScript IV First-Strand Synthesis System with random hexamers (Invitrogen) and subsequently stored at −20°C. Specific primers used in this study are listed in Table S1 and were obtained from a commercial supplier (Microsynth, Switzerland). qRT-PCR reactions were prepared with SYBRGreen PCR Master Mix (Applied Biosystems) with 1 μM primers, and 1 or 2 μL cDNA. Reactions were run as absolute quantification on ABI PRISM7900HT Sequence Detection System (Applied Biosystems) or QuantStudio 7 system (Applied Biosystems). Amplicon specificity was confirmed by melting-curve analysis.

### Statistical analysis

Statistical analyses are described in each figure legend. For experiments combining several groups, one-way ANOVA with Dunn’s multiple comparison test was used. For comparisons between two experimental groups, the two-tailed Mann-Whitney or Welch’s test was used. Statistical significance was determined using Prism v.9.5.0 software (GraphPad). P > 0.05 was considered non-significant. For time-lapse image analysis, fields of view from a given LoC were considered as biological replicates. All experiments were repeated at least twice.

## References

1. Weibel, E.R. (2015). On the Tricks Alveolar Epithelial Cells Play to Make a Good Lung. Am J Respir Crit Care Med 191, 504–513. 10.1164/rccm.201409-1663OE.

2. Notter, R.H. (2000). Lung surfactants: Basic science and clinical applications. Lung Surfactants: Basic Science and Clinical Applications 149, 1–444. 10.1201/9781482270426/.

3. Goerke, J. (1998). Pulmonary surfactant: Functions and molecular composition. Preprint at Elsevier, 10.1016/S0925-4439(98)00060-X 10.1016/S0925-4439(98)00060-X.

4. Whitsett, J.A., and Weaver, T.E. (2002). Hydrophobic Surfactant Proteins in Lung Function and Disease. New England Journal of Medicine 347, 2141–2148. 10.1056/NEJMra022387.

5. Clements, J.A., Nellenbogen, J., and Trahan, H.J. (1970). Pulmonary Surfactant and Evolution of the Lungs. Science (1979) 169, 603–604. 10.1126/SCIENCE.169.3945.603.

6. Hall, S.B., and Zuo, Y.Y. (2024). The biophysical function of pulmonary surfactant. Biophys J 123, 1519–1530. 10.1016/J.BPJ.2024.04.021/.

7. Torrelles, J.B., and Schlesinger, L.S. (2017). Integrating Lung Physiology, Immunology, and Tuberculosis. Trends Microbiol 25, 688–697. 10.1016/J.TIM.2017.03.007.

8. Van Iwaarden, J.F., Claassen, E., Jeurissen, S.H.M., Haagsman, H.P., and Kraal, G. (2012). Alveolar Macrophages, Surfactant Lipids, and Surfactant Protein B Regulate the Induction of Immune Responses via the Airways. 10.1165/ajrcmb.24.4.4239 24, 452–458. 10.1165/AJRCMB.24.4.4239.

9. Andreeva, A. V., Kutuzov, M.A., and Voyno-Yasenetskaya, T.A. (2007). Regulation of surfactant secretion in alveolar type II cells. American Journal of Physiology-Lung Cellular and Molecular Physiology 293, L259–L271. 10.1152/ajplung.00112.2007.

10. Hidalgo, A., Garcia-Mouton, C., Autilio, C., Carravilla, P., Orellana, G., Islam, M.N., Bhattacharya, J., Bhattacharya, S., Cruz, A., and Pérez-Gil, J. (2021). Pulmonary surfactant and drug delivery: Vehiculization, release and targeting of surfactant/tacrolimus formulations. Journal of Controlled Release 329, 205–222. 10.1016/J.JCONREL.2020.11.042.

11. Wang, J., Li, P., Yu, Y., Fu, Y., Jiang, H., Lu, M., Sun, Z., Jiang, S., Lu, L., and Wu, M.X. (2020). Pulmonary surfactant–biomimetic nanoparticles potentiate heterosubtypic influenza immunity. Science (1979) 367, eaau0810. 10.1126/science.aau0810.

12. Katsura, H., Sontake, V., Tata, A., Kobayashi, Y., Edwards, C.E., Heaton, B.E., Konkimalla, A., Asakura, T., Mikami, Y., Fritch, E.J., et al. (2020). Human Lung Stem Cell-Based Alveolospheres Provide Insights into SARS-CoV-2-Mediated Interferon Responses and Pneumocyte Dysfunction. Cell Stem Cell 27, 890–904.e8. 10.1016/J.STEM.2020.10.005.

13. Di Domizio, J., Gulen, M.F., Saidoune, F., Thacker, V. V., Yatim, A., Sharma, K., Nass, T., Guenova, E., Schaller, M., Conrad, C., et al. (2022). The cGAS-STING pathway drives type I IFN immunopathology in COVID-19. Nature 2022, 1–9. 10.1038/s41586-022-04421-w.

14. Deinhardt-Emmer, S., Rennert, K., Schicke, E., Cseresnyés, Z., Windolph, M., Nietzsche, S., Heller, R., Siwczak, F., Haupt, K.F., Carlstedt, S., et al. (2020). Co-infection with Staphylococcus aureus after primary influenza virus infection leads to damage of the endothelium in a human alveolus-on-a-chip model. Biofabrication 12, 025012. 10.1088/1758-5090/ab7073.

15. Hoang, T.N.M., Cseresnyés, Z., Hartung, S., Blickensdorf, M., Saffer, C., Rennert, K., Mosig, A.S., von Lilienfeld-Toal, M., and Figge, M.T. (2022). Invasive aspergillosis-on-chip: A quantitative treatment study of human Aspergillus fumigatus infection. Biomaterials 283, 121420. 10.1016/J.BIOMATERIALS.2022.121420.

16. Felder, M., Stucki, A.O., Stucki, J.D., Geiser, T., and Guenat, O.T. (2014). The potential of microfluidic lung epithelial wounding: towards in vivo-like alveolar microinjuries. Integr. Biol. 6, 1132–1140. 10.1039/C4IB00149D.

17. Zamprogno, P., Wüthrich, S., Achenbach, S., Thoma, G., Stucki, J.D., Hobi, N., Schneider-Daum, N., Lehr, C.M., Huwer, H., Geiser, T., et al. (2021). Second-generation lung-on-a-chip with an array of stretchable alveoli made with a biological membrane. Communications Biology 2021 4:1 4, 168-. 10.1038/s42003-021-01695-0.

18. Huh, D., Matthews, B.D., Mammoto, A., Montoya-Zavala, M., Hsin, H.Y., and Ingber, D.E. (2010). Reconstituting organ-level lung functions on a chip. Science 328, 1662–1668. 10.1126/science.1188302.

19. Stucki, A.O., Stucki, J.D., Hall, S.R.R., Felder, M., Mermoud, Y., Schmid, R.A., Geiser, T., and Guenat, O.T. (2015). A lung-on-a-chip array with an integrated bio-inspired respiration mechanism. Lab Chip 15, 1302–1310. 10.1039/c4lc01252f.

20. Thacker, V. V., Dhar, N., Sharma, K., Barrile, R., Karalis, K., and McKinney, J.D. (2020). A lung-on-chip model of early M. Tuberculosis infection reveals an essential role for alveolar epithelial cells in controlling bacterial growth. Elife 9, 1–73. 10.7554/eLife.59961.

21. Paraskeva, M.A., Westall, G.P., and Snell, G.I. (2020). Immunosuppressive strategies in lung transplantation. Ann Transl Med 8, 409. 10.21037/ATM.2019.12.117.

22. Deuse, T., Blankenberg, F., Haddad, M., Reichenspurner, H., Phillips, N., Robbins, R.C., and Schrepfer, S. (2012). Mechanisms behind Local Immunosuppression Using Inhaled Tacrolimus in Preclinical Models of Lung Transplantation. Am J Respir Cell Mol Biol 43, 403–412. 10.1165/RCMB.2009-0208OC.

23. Hidalgo, A., Salomone, F., Fresno, N., Orellana, G., Cruz, A., and Perez-Gil, J. (2017). Efficient Interfacially Driven Vehiculization of Corticosteroids by Pulmonary Surfactant. Langmuir 33, 7929–7939. 10.1021/ACS.LANGMUIR.7B01177/.

24. Misharin, A. V., Morales-Nebreda, L., Reyfman, P.A., Cuda, C.M., Walter, J.M., McQuattie-Pimentel, A.C., Chen, C.I., Anekalla, K.R., Joshi, N., Williams, K.J.N., et al. (2017). Monocyte-derived alveolar macrophages drive lung fibrosis and persist in the lung over the life span. Journal of Experimental Medicine 214, 2387–2404. 10.1084/jem.20162152.

25. Pahari, S., Arnett, E., Simper, J., Azad, A., Guerrero-Arguero, I., Ye, C., Zhang, H., Cai, H., Wang, Y., Lai, Z., et al. (2023). A new tractable method for generating human alveolar macrophage-like cells in vitro to study lung inflammatory processes and diseases. mBio 14, e0083423. 10.1128/MBIO.00834-23/.

26. Jacob, A., Morley, M., Hawkins, F., McCauley, K.B., Jean, J.C., Heins, H., Na, C.-L., Weaver, T.E., Vedaie, M., Hurley, K., et al. (2017). Differentiation of Human Pluripotent Stem Cells into Functional Lung Alveolar Epithelial Cells. Cell Stem Cell 21, 472–488.e10. 10.1016/j.stem.2017.08.014.

27. Yamamoto, Y., Gotoh, S., Korogi, Y., Seki, M., Konishi, S., Ikeo, S., Sone, N., Nagasaki, T., Matsumoto, H., Muro, S., et al. (2017). Long-term expansion of alveolar stem cells derived from human iPS cells in organoids. Nat Methods 14, 1097–1106. 10.1038/nmeth.4448.

28. Guilliams, M., De Kleer, I., Henri, S., Post, S., Vanhoutte, L., De Prijck, S., Deswarte, K., Malissen, B., Hammad, H., and Lambrecht, B.N. (2013). Alveolar macrophages develop from fetal monocytes that differentiate into long-lived cells in the first week of life via GM-CSF. Journal of Experimental Medicine 210, 1977–1992. 10.1084/JEM.20131199.

29. Yu, X., Buttgereit, A., Lelios, I., Utz, S.G., Cansever, D., Becher, B., and Greter, M. (2017). The Cytokine TGF-β Promotes the Development and Homeostasis of Alveolar Macrophages. Immunity 47, 903–912.e4. 10.1016/j.immuni.2017.10.007.

30. Vogt, M.T., Thomas, C., Vassallo, C.L., Basford, R.E., and Gee, J.B. (1971). Glutathione-dependent peroxidative metabolism in the alveolar macrophage. J Clin Invest 50, 401–410. 10.1172/JCI106507.

31. Rothchild, A.C., Olson, G.S., Nemeth, J., Amon, L.M., Mai, D., Gold, E.S., Diercks, A.H., and Aderem, A. (2019). Alveolar macrophages generate a noncanonical NRF2-driven transcriptional response to Mycobacterium tuberculosis in vivo. Sci Immunol 4. 10.1126/sciimmunol.aaw6693.

32. Dodd, C.E., Pyle, C.J., Glowinski, R., Rajaram, M.V.S., and Schlesinger, L.S. (2016). CD36-Mediated Uptake of Surfactant Lipids by Human Macrophages Promotes Intracellular Growth of Mycobacterium tuberculosis. The Journal of Immunology 197, 4727–4735. 10.4049/jimmunol.1600856.

33. Mishra, R., Hannebelle, M., Patil, V.P., Dubois, A., Garcia-Mouton, C., Kirsch, G.M., Jan, M., Sharma, K., Guex, N., Sordet-Dessimoz, J., et al. (2023). Mechanopathology of biofilm-like Mycobacterium tuberculosis cords. Cell 186, 5135–5150.e28. 10.1016/j.cell.2023.09.016.

34. Thacker, V. V, Sharma, K., Dhar, N., Mancini, G., Sordet-Dessimoz, J., and McKinney, J.D. (2021). Rapid endotheliitis and vascular damage characterize SARS-CoV-2 infection in a human lung-on-chip model. EMBO Rep 22. 10.15252/embr.202152744.

35. Olivier, K.N., Shaw, P.A., Glaser, T.S., Bhattacharyya, D., Fleshner, M., Brewer, C.C., Zalewski, C.K., Folio, L.R., Siegelman, J.R., Shallom, S., et al. (2014). Inhaled amikacin for treatment of refractory pulmonary nontuberculous mycobacterial disease. Ann Am Thorac Soc 11, 30–35. 10.1513/ANNALSATS.201307-231OC/.

36. Pearce, C., Ruth, M.M., Pennings, L.J., Wertheim, H.F.L., Walz, A., Hoefsloot, W., Ruesen, C., Muñoz Gutiérrez, J., Gonzalez-Juarrero, M., and Van Ingen, J. (2020). Inhaled tigecycline is effective against Mycobacterium abscessus in vitro and in vivo. Journal of Antimicrobial Chemotherapy 75, 1889. 10.1093/JAC/DKAA110.

37. Dartois, V., and Dick, T. (2024). Therapeutic developments for tuberculosis and nontuberculous mycobacterial lung disease. Nature Reviews Drug Discovery 2024 23:5 23, 381–403. 10.1038/s41573-024-00897-5.

38. Dartois, V., and Dick, T. (2024). Toward better cures for Mycobacterium abscessus lung disease. Clin Microbiol Rev, e0008023. 10.1128/CMR.00080-23.

39. van Heeswijk, R.P.G., Dannemann, B., and Hoetelmans, R.M.W. (2014). Bedaquiline: a review of human pharmacokinetics and drug-drug interactions. Journal of Antimicrobial Chemotherapy 69, 2310–2318. 10.1093/jac/dku171.

40. Labuda, S.M., Seaworth, B., Dasgupta, S., and Goswami, N.D. (2024). Bedaquiline, pretomanid, and linezolid with or without moxifloxacin for tuberculosis. Lancet Respir Med 12, e5–e6. 10.1016/S2213-2600(23)00426-5.

41. Ruth, M.M., Sangen, J.J.N., Remmers, K., Pennings, L.J., Svensson, E., Aarnoutse, R.E., Zweijpfenning, S.M.H., Hoefsloot, W., Kuipers, S., Magis-Escurra, C., et al. (2019). A bedaquiline/clofazimine combination regimen might add activity to the treatment of clinically relevant non-tuberculous mycobacteria. J Antimicrob Chemother 74, 935–943. 10.1093/JAC/DKY526.

42. García-Mouton, C., Echaide, M., Serrano, L.A., Orellana, G., Salomone, F., Ricci, F., Pioselli, B., Amidani, D., Cruz, A., and Pérez-Gil, J. (2023). Beyond the Interface: Improved Pulmonary Surfactant-Assisted Drug Delivery through Surface-Associated Structures. Pharmaceutics 15, 256. 10.3390/PHARMACEUTICS15010256/S1.

43. Collada, A., Cruz, A., and Pérez-Gil, J. (2025). Studying the interfacial activity and structure of pulmonary surfactant complexes. Chem Phys Lipids 266, 105459. 10.1016/J.CHEMPHYSLIP.2024.105459.

44. Burgess, C.L., Ayers, L.J., Minakin, K., Alysandratos, K.D., Varelas, X., and Kotton, D.N. (2025). Protocol for the differentiation of human alveolar epithelial type I cells from pluripotent stem cell-derived type II-like cells. STAR Protoc 6, 103667. 10.1016/J.XPRO.2025.103667.

45. Dobin, A., Davis, C.A., Schlesinger, F., Drenkow, J., Zaleski, C., Jha, S., Batut, P., Chaisson, M., and Gingeras, T.R. (2013). STAR: ultrafast universal RNA-seq aligner. Bioinformatics 29, 15–21. 10.1093/BIOINFORMATICS/BTS635.

46. Chen, Y., Lun, A.T.L., Smyth, G.K., Burden, C.J., Ryan, D.P., Khang, T.F., and Lianoglou, S. (2016). From reads to genes to pathways: differential expression analysis of RNA-Seq experiments using Rsubread and the edgeR quasi-likelihood pipeline. F1000Research 2016 5:1438 5, 1438. 10.12688/f1000research.8987.2.

47. Thacker, V. V, Dhar, N., Sharma, K., Barrile, R., Karalis, K., and McKinney, J.D. (2020). A lung-on-chip model of early M. tuberculosis infection reveals an essential role for alveolar epithelial cells in controlling bacterial growth. Elife 9. 10.7554/elife.59961.

48. Thacker, V. V, Sharma, K., Dhar, N., Mancini, G., Sordet-Dessimoz, J., and McKinney, J.D. (2021). Rapid endotheliitis and vascular damage characterize SARS-CoV-2 infection in a human lung-on-chip model. EMBO Rep 22. 10.15252/embr.202152744.

49. Taeusch, H.W., De La Serna, J.B., Perez-Gil, J., Alonso, C., and Zasadzinski, J.A. (2005). Inactivation of pulmonary surfactant due to serum-inhibited adsorption and reversal by hydrophilic polymers: Experimental. Biophys J 89, 1769–1779. 10.1529/BIOPHYSJ.105.062620/.

50. Schürch, D., Ospina, O.L., Cruz, A., and Pérez-Gil, J. (2010). Combined and Independent Action of Proteins SP-B and SP-C in the Surface Behavior and Mechanical Stability of Pulmonary Surfactant Films. Biophys J 99, 3290–3299. 10.1016/J.BPJ.2010.09.039.

