## Supplementary Note 1 for "Vehiculation and functional delivery of lipophilic therapeutics and antibiotics via pulmonary surfactant in a lung-on-chip model"

### ***Surfactant formulations***

| Sample | Concentration | Solvent |
|---|---|---|
| Organic Extract from Native Surfactant (OE) | 60.1 mg/mL (phospholipids) | Chloroform:Methanol |
| DPPC | 41.3 mg/mL | Chloroform:Methanol |
| POPG | 50 mg/mL | Chloroform:Methanol |
| RhodDOPE [Avanti, #810150P] | 1 mg/mL | Methanol |
| TacNB | 1.7 mg/mL | Chloroform:Methanol |
| Tacrolimus | 10 mg/mL | Chlo:MetOH |
| Bedaquiline | 10 mg/mL | DMSO |
| 18:1 Cy5 PC | 250 μg/mL | Chloroform |

To calculate the % mol/mol, we used the molecular weight of DPPC (734,04 g/mol) as the MW of surfactant.

**OE-RhodDOPE 1% mol/mol + TacNB 2% w/w:**

Liquid administration:

1. 4.16 μL OE + 4.43 μL RhodDOPE + 3 μL TacNB
2. Dry and reconstitute with 50 μL PBS. Lipid concentration is 5 mg/mL.
3. Dilute to 2 mg/mL lipid concentration in alveolar epithelial cell media.

TFB administration:

1. 24.96 μL OE + 26.4 μL RhodDOPE + 17.7 μL TACNB
2. Dry and reconstitute with 30 μL PBS. Lipid concentration is 50 mg/mL.

**MLVs (DPPC:POPG 7:3 w/w) + RhodDOPE 1% mol/mol + TacNB 2% w/w:**

Liquid administration:

1. 4.24 μL DPPC + 1.5 μL POPG + 4.43 μL RhodDOPE + 3 μL TacNB
2. Dry and reconstitute with 50 μL PBS. Lipid concentration is 5 mg/mL.
3. Dilute to 2 mg/mL lipid concentration in alveolar epithelial cell media.

**OE- Cy5-PC 1% mol/mol + Bedaquiline 2% w/w:**

Bridge administration:

1. 16.64 μL OE + 71.68 μL of Cy5-PC
2. Dry and incubate for 30 min at 45 ºC in 18 μL PBS
3. Add 2 μL Bedaquiline (10 mg/mL in DMSO) and incubate for 30 min at 45 ºC.
